# Preconditioning cathodal transcranial direct current stimulation facilitates the neuroplastic effect of subsequent anodal transcranial direct current stimulation applied during cycling in young adults

**DOI:** 10.1101/653634

**Authors:** Maryam Pourmajidian, Benedikt Lauber, Simranjit K Sidhu

## Abstract

The study aimed to examine the effect of a priming cathodal transcranial direct current stimulation (ctDCS) before subsequent anodal-tDCS (atDCS) was applied during low workload cycling exercise on the corticospinal responses in young healthy individuals. Eleven young subjects participated in two sessions receiving either priming ctDCS or sham stimulation, followed by atDCS while cycling (i.e. ctDCS-atDCS, sham-atDCS) at 1.2 times their body weight (84 ± 20 W) in a counterbalanced double-blind design. Corticospinal excitability was measured with motor evoked potentials (MEPs) elicited via transcranial magnetic stimulation with the intensity set to produce an MEP amplitude of 1 mV in a resting hand muscle at baseline (PRE), following priming tDCS (POST-PRIMING) and post atDCS combined with cycling exercise (POST-TEST). There was a significant interaction between time and intervention (*P* < *0.01*) on MEPs. MEPs increased from PRE (1.0 ± 0.06 mV) to POST-TEST (1.3 ± 0.06 mV) during ctDCS-atDCS (*P* < *0.001*) but did not change across time during sham-atDCS (1.0 ± 0.06 mV, *P* > *0.7*). Furthermore, MEPs were higher in ctDCS-atDCS compared to sham-atDCS (*P* < *0.01*) at both POST-PRIMING (ctDCS-atDCS: 1.1 ± 0.06, sham-atDCS: 1.0 ± 0.06) and POST-TEST (ctDCS-atDCS: 1.3 ± 0.06, sham-atDCS: 1.0 ± 0.06). These outcomes demonstrate that cathodal tDCS priming can enhance corticospinal excitability following anodal tDCS applied in combination with cycling exercise. The findings have implications for the application of tDCS in combination with cycling exercise in rehabilitation and sporting contexts.

## Introduction

It is relatively well established that regular participation in whole body exercise can have a positive impact on intrinsic brain network plasticity and connectivity [5, 13]; and translate into functional behavioral improvements in cognitive and motor performance [19, 22]. Despite these findings, studies probing the influence of whole body exercise on corticospinal excitability provide mixed outcomes. For example, low intensity cycling exercise does not modulate net corticospinal tract excitability, but attenuates intracortical inhibition [23, 39, 41]. It is speculated that the attenuation in intracortical inhibition likely contributes to the creation of a cortical environment that is optimal for plasticity. However, the lack of a change in corticospinal excitability makes this interpretation difficult. It is possible that the effect of cycling exercise on corticospinal excitably represents a homeostatic metaplastic mechanism, although, this remains elusive [4].

Non-invasive brain stimulation (NIBS) is known to induce neuroplastic changes in the brain. These modifications are manifested in either long-term potentiation (LTP) or long-term depression (LTD) of synapses in favour of improving motor and cognitive function in healthy adults and individuals with neurological impairments [25]. Transcranial direct current stimulation (tDCS) is an emerging NIB S technique which can be used to deliver weak electrical currents to the brain in order to modulate the excitability of the primary motor cortex (M1) in humans [25]. Cathodal tDCS (i.e. ctDCS; cathode placed over M1) is known to produce LTD-like effects while anodal tDCS (i.e. atDCS; anode placed over M1) has been used to elicit LTP-like effects [26]. A few recent studies have demonstrated the effectiveness of atDCS in modulating corticospinal excitability and motor performance [1, 3, 10, 45]. However, the outcomes in relation to its efficacy are variable, likely represented via differences in experimental manipulations, such as site, intensity and duration of stimulation, as well as inter-individual variability [15, 20, 43, 48].

There is increasing evidence to demonstrate that priming the brain by reducing the activation threshold of the neurons can augment the effect of subsequent stimulation inducing larger LTP-like effect [14, 17, 38]. A possible mechanism to explain the enhancing effect of priming stimulation is homeostatic metaplasticity [6], which proposes that synaptic plasticity is bidirectional and the previous level of synaptic activity may alter the response to the subsequent NIBS. Essentially, higher post synaptic activity will increase the synaptic modification threshold, while lower post synaptic activity will decrease the threshold and facilitate the induction of LTP-like effects [6]. In agreement with this proposition, studies using tDCS have demonstrated that priming cathodal tDCS can amplify the effect of subsequent anodal tDCS to increase corticospinal excitability and improve motor skill performance [8, 12]. For instance, when cathodal tDCS precedes anodal tDCS, completion time of a grooved pegboard test is significantly reduced compared to shamtDCS – atDCS and shamtDCS – shamtDCS conditions [8].

The aim of the current study was to investigate the effects of priming ctDCS on corticospinal excitability tested with transcranial magnetic stimulation (TMS) evoked potential (MEP) after low intensity cycling exercise performed with concurrent atDCS in young healthy individuals. We hypothesised that, priming ctDCS (but not priming sham) will enhance the LTP-like effects of the subsequent atDCS combined with a low intensity cycling exercise to augment corticospinal excitability.

## Methods

### Subjects

Eleven young (20.9 ± 0.2 years; 4 females) healthy subjects (height: 172 ± 2cm; weight: 71 ± 3kg) were recruited through advertisement in the university. All subjects were right handed (handedness laterality index: 0.86 ± 0.02) in accordance with the Edinburgh Handedness Inventory [27]. Participants with contraindications to TMS or tDCS including a history of epilepsy, stroke, neurological illness, or those who were consuming psychoactive medications at the time of the study were excluded from participation. Each subject gave their written informed consent prior to participation and was instructed to avoid any strenuous activity at least 24 hrs prior to the experimental sessions. The study was approved by the University of Adelaide Human Research Ethics Committee and conducted in accordance with the Declaration of Helsinki.

### Experimental Protocol

Subjects were set-up on the cycle ergometer with their feet strapped into the pedals and were asked to minimize the motion of their right wrist, forearm and hand throughout the experiment. There were two experimental sessions, and each session was separated by a week. In addition, both sessions were conducted between 2 pm to 5 pm and repeat sessions were conducted at the same time of the day to minimise the confounding influence of diurnal variations in cortisol on cortical plasticity [33].

During the two sessions, subjects received a dose of priming tDCS of either sham or ctDCS in a counter-balanced, double-blinded approach. Subjects then performed a low intensity cycling exercise on a mechanically braked cycle ergometer (Velotron, Elite Model, Racer Mate, Seattle, WA) for 10 minutes (power output of the cycle ergometer was set at 120 percent of participant’s body weight; group mean workload = 85 ± 3 Watts; and 80 rpm fixed cadence) while atDCS was applied over the M1 area concurrently. The rationale for implementing this cycling intensity (i.e. < 100 W) was so that the participants will be subjected to an intensity of exercise, that did not induce fatigue (whilst corrected for their body weight) – which can independently influence corticospinal tract excitability [36, 37].

MEPs (i.e. 3 blocks of 15 TMS) were collected over a 10 min period at three time points; baseline, immediately post priming stimulation and immediately post atDCS combined with cycling exercise (i.e. PRE, POST-PRIMING and POST-TEST respectively) (Fig. 1).

**Figure 1.**
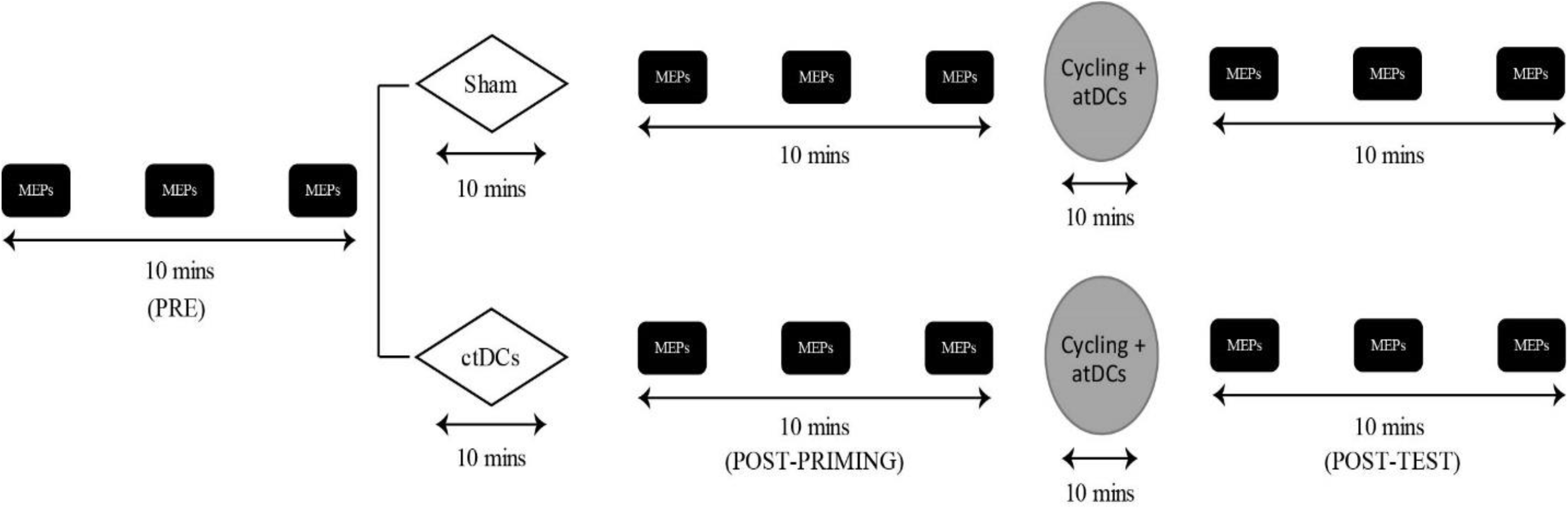
Schematic of experimental protocol. Resting motor threshold and stimulus intensity to evoke a 1mV response was measured at the start of the experimental session (not shown in figure). A set of baseline MEPs (i.e. 3 blocks of 15 transcranial magnetic stimulations) was collected in both sessions as baseline (PRE) prior to priming tDCS. Subjects were given a priming dose of either ctDCS (10 min at 2 mA) or sham (30 s stimulation + 9.5 min no stimulation) immediately after PRE. MEP measurements were repeated immediately after priming tDCS (POST-PRIMING) and atDCS combined with cycling exercise (POST-TEST).

### Experimental Procedures

#### Electromyography recordings

After skin preparation with abrasion and alcohol swabs, surface electrodes were positioned over the muscle belly and tendon of the right first dorsal interosseous (FDI). EMG was recorded via a monopolar configuration (Ag-AgCl, 10 mm diameter, inter-electrode distance: 2.9 ± 0.1 cm). EMG signals were amplified (100-1000 times; MA300 DTU, Motion Lab Systems, USA), band pass filtered (30-1000 Hz) and analogue to digitally converted at a sampling rate of 2000 Hz using a 16-bit power 1401 and Signal 4.11 data collection software (Cambridge Electronic Design, UK) via custom written scripts. Collected data was stored on a laboratory PC for offline analysis.

#### Transcranial magnetic stimulation (TMS)

Single-pulse TMS was delivered to the left M1 using a MagStim 200 (monophasic) magnetic stimulator (MagStim, Dyfed, UK) and a figure-of-eight coil. The coil was placed tangentially to the scalp with an angle of approximately 45° posteriorly producing a current flow within the motor cortex in the posterior-anterior direction. The optimal location that gave the largest MEP at a fixed stimulation intensity in the resting FDI hand muscle was marked directly on the scalp to ensure consistent positioning of the coil. Resting motor threshold (RMT) was defined as the minimum stimulator intensity (MSO) which produced an MEP amplitude of at least 50μv in 5 out of 10 successive stimulations was determined to ensure that the effective TMS intensity set to produce an MEP peak-to-peak amplitude of 1 mV in the resting FDI was consistent across sessions.

#### Transcranial direct current stimulation

tDCS was delivered through a direct current electrical stimulator (NeuroConn DC-Stimulator, Germany) connected to a pair of saline soaked electrodes (25 cm^2^). For ctDCS, the cathode was placed over the left motor cortex with anode on the right supraorbital area and for atDCS, the placement of the electrodes was reversed [46]. Stimulation was applied at an intensity of 2 mA for a period of 10 min (atDCS, ctDCS) or 30 s followed by 9.5 mins of no stimulation (sham) [48]. In order to minimise the discomfort caused by electrical transients, 10 s of fade in and fade out current was set at the start and end of the stimulation period [25]. The selection of priming tDCS (either ctDCS or sham) and test tDCS (atDCS) in each of the two experimental sessions was double-blinded to both the subject and the main experimenter. At the end of each tDCS dose, subjects were asked to report the sensations associated with the stimulation. It was clear from the subjects’ subjective responses that the type of stimulation received was not discernible and subjects mainly felt slight tingling at the site of the electrode [47].

### Data Analysis

Signal 4.11 software was used for offline analysis of the EMG data. Since the aim was to quantify corticospinal excitability (CSE) in a resting FDI muscle, in trials where muscle activity (measured as peak to peak EMG) was more than 20 μV in the 100 ms prior to stimulation, the data was removed from the analysis (< 1% of trials). MEP amplitude was measured as peak-to-peak amplitude and expressed in mV.

### Statistical Analysis

Normality of the data was confirmed by a Shapiro-Wilk W test. One-way repeated measures analysis of variance (ANOVA) were used to compare RMT and MEP amplitude across sessions (i.e. sham-atDCS; ctDCS-atDCS). Linear mixed model analyses with repeated measures were used to investigate the effect of time (i.e. PRE, POST-PRIMING and POST-TEST) and intervention (i.e. sham-atDCS, ctDCS-atDCS) on MEP amplitude. For linear mixed model analyses, subjects were included as random effect, and significant main effects and interactions were further investigated using custom contrasts with Bonferroni correction. Cohen’s effect sizes were calculated with G * Power software. Data (in text and figures) are presented as mean ± standard error of the mean. Statistical significance is set at *P* < 0.05.

## Results

All participants completed the study with no adverse reaction. There was no difference in RMT (41.9 ± 0.7 % MSO for sham-atDCS and 40.9 ± 0.9 % MSO for ctDCS-atDCS) and S_1mv_ (53.3 ± 1.4 % MSO for sham-atDCS and 51.1 ± 1.4 % MSO for ctDCS-atDCS) between interventions (*F_1,21_* < *0.4, P* > *0.5*, d_Z_ < 0.13).

MEP amplitude (Fig. 2) differed as a function of time (*F_2, 774_ = 4.7, P* < 0.01; d_Z_ = 0.95) and intervention (*F_1, 303_ = 21.1, P* < 0.001; d_Z_ = 1.45). In addition, there was an interaction between factors (*F_2, 787_= 5.9, P* < *0.01*; d_Z_ = 1.75). MEP amplitude did not change from PRE to POST-PRIMING (*P* = *0.07*, d_Z_ = 1.08), however, it significantly increased from PRE to POST-TEST (*P* < *0.001*, d_Z_ = 2.13) in ctDCS-atDCS session. While MEP amplitude did not modulate across time in sham-atDCS session (*P* > 0.9, d_Z_ < 0.28), it was significantly lower in sham-atDCS session compared to ctDCS-atDCS session at both POST-PRIMING and POST-TEST time points (*P* < *0.01*, d_Z_ > 1.42).

**Figure 2.**
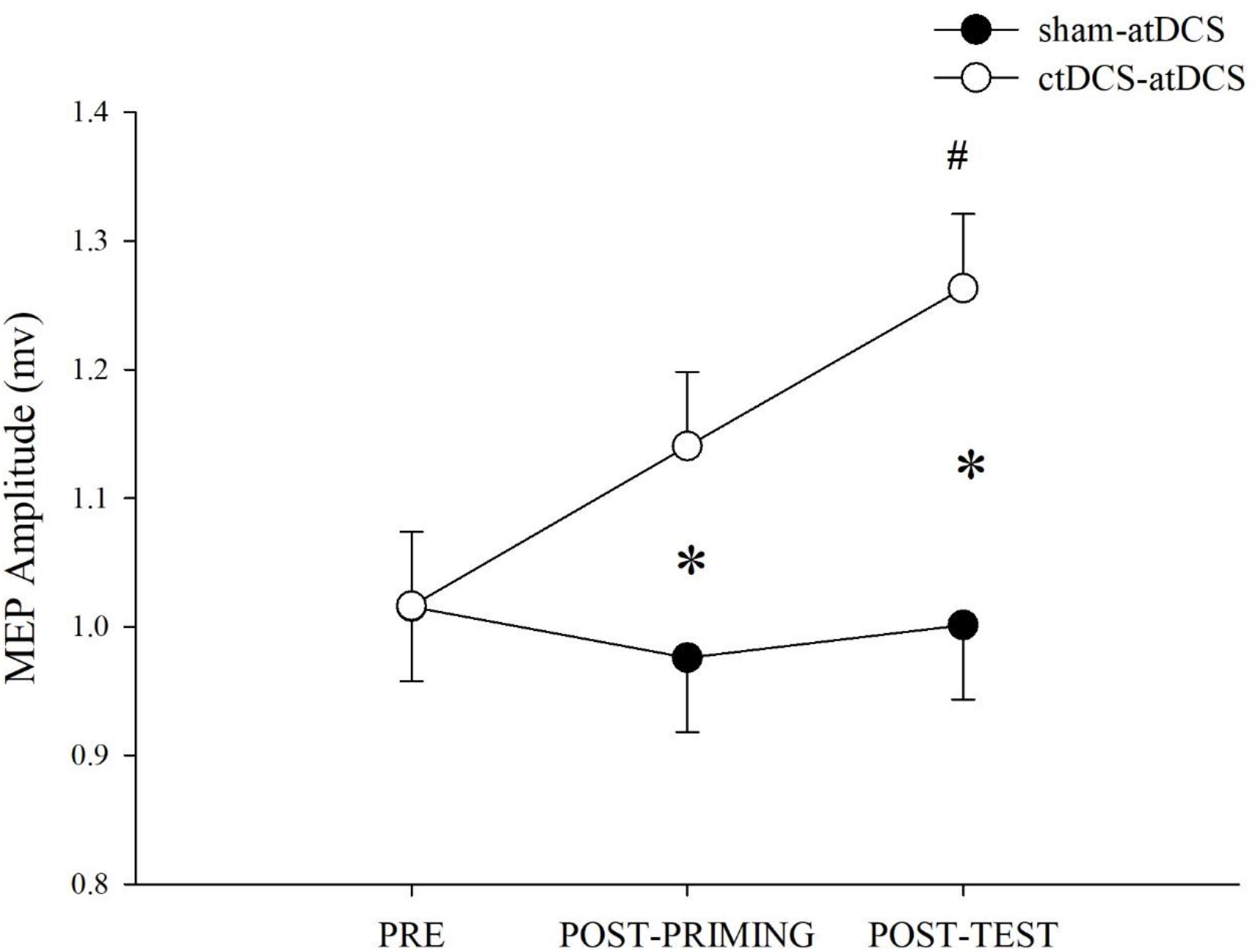
Corticospinal excitability (i.e. MEP amplitude) at baseline (PRE), after ctDCS or sham priming (POST-PRIMING) and after cycling exercise performed with atDCS (POST-TEST) in the two sessions. Data are expressed as the MEP amplitude (Mean ± SE). \**P*<0.05 between sessions; ^#^*P*<0.05 from PRE in ctDCS - atDCS session.

## Discussion

The present study aimed to investigate whether the efficacy of anodal tDCS applied during low workload cycling exercise over M1 to modulate corticospinal excitability will alter when primed by cathodal tDCS applied over M1. The main outcome of the study was that priming cathodal tDCS augmented corticospinal tract excitability following anodal tDCS applied in combination with cycling exercise. Specifically, MEPs increased (by approximately thirty percent) compared to baseline when anodal tDCS applied during cycling exercise was primed with cathodal tDCS; compared to when preconditioned using sham tDCS. This finding suggests that ctDCS priming may potentially be used as a tool to improve the neuroplastic response to cycling exercise and consequently motor/cognitive function.

### tDCS mediated effect on the neuroplastic response to cycling exercise

There is increasing evidence that locomotor exercise create a cortical environment that is optimal for neuroplasticity [9, 11], and may consequently have a positive influence on brain structure and functions, including memory and motor skill learning [22, 40]. The underlying neurophysiological mechanisms of the neuroplastic response to locomotor cycling remain elusive, although animal work suggest that increased levels of BDNF may contribute [44]. Interestingly, even though human studies using non-invasive brain stimulation techniques suggest that a short bout of cycling exercise attenuates the magnitude of GABA_A_ – mediated inhibition [39, 41], these studies have shown no change in corticospinal excitability measured with a TMS-evoked MEP response [23, 39, 41]. The mechanism underlying the lack of a change in the corticospinal tract excitability post cycling exercise remains unknown. However, it is well-understood that the induction of neuroplastic response in healthy humans is influenced by many factors, including the history of activity within the targeted neuronal network [18, 32] which is thought to be particularly important. For example, synaptic potentiation is dampened following high neuronal activity (e.g. repetitive locomotor movements that are known to increase cortical excitability [31, 34]), but increased following low neural activity [2, 16]; an effect that represents homeostatic metaplasticity to maintain network integrity [2]. Therefore, it is possible that the lack of a neuroplastic response (i.e. no change in an MEP) reported post single bout of cycling exercise in previous work [23, 39, 41] is in accordance with Bienenstock-Cooper-Munro theory of homeostatic metaplasticity [6]. Our study provides an important and novel finding to support this notion. We used tDCS to induce activity-dependent plasticity (i.e. long-term depression; LTD and long-term potentiation; LTP) at pre-and-during low workload cycling exercise and show a homeostatic metaplastic response in the individuals studied. This experimental design is prudent since contrary to other forms of neuromodulation (e.g. paired associative stimulation and repetitive TMS [21, 30], tDCS can be applied simultaneously with motor activity to influence corticospinal excitability and promote function [7, 42]. Specifically, MEPs were augmented post-cycling exercise when ctDCS (which is known to induce LTD-like effect by decreasing the neuronal activity to reduce the threshold for subsequent stimulations that increase excitability of neurons [38]) was used to prime the atDCS applied during cycling exercise. This effect was not apparent when priming sham stimulation was applied. The outcomes of this study provide important evidence to demonstrate that lowering the threshold for LTP-like induction may in fact be necessary to induce LTP-like plasticity with atDCS during cycling exercise.

The priming effect observed in the current study with ctDCS over M1 (i.e. no significant change – but tendency for an increase) is consistent with the findings in a recent study where no change [12] or even an increase [4] was evident after 10 – 20 minutes of priming ctDCS. The variability in the response to tDCS protocols may partly be explained by the fact that about 50% of the population have poor or absent responses to tDCS [48]. It is however important to note that priming stimulation can still be effective at inducing metaplasticity even without modulation of synaptic efficacy [2] or an overt modulation of MEP amplitude [7, 28, 29, 35]. For example, it has been shown that priming with NIBS that induces LTP or LTD did not alter MEPs as standalone, but in fact had an opposing effect on subsequent NIBS response in young and older subjects [28, 35].

### Methodological considerations

There are several limitations in the present study. First, we did not measure corticospinal excitability after cycling exercise without the application of tDCS. However, the lack of a modulation in corticospinal excitability post normal cycling exercise has been documented in numerous previous studies in both exercised and non-exercised muscles [23, 39, 41]; and the fact that we did not observe any change post shamtDCS-atDCS session suggests that it is unlikely a change would have been observed after cycling (without concurrent atDCS). In any case, the objective of the study was to establish an optimal tDCS priming paradigm applied during cycling that would augment the corticospinal excitability. Another consideration is the fact that the development of homeostatic metaplasticity is dependent on the timing between priming and subsequent neuromodulatory stimulation, whereby a longer interval of 30-min has recently been indicated to be most effective [24, 35]. While the priming and test neuromodulation were separated by 10 minutes in the current study, future studies should probe the effect of time in between priming tDCS and atDCS applied during cycling exercise on the activity-dependant metaplastic response. The small sample size of this study should be acknowledged; however, it should also be noted that the effect sizes for all significant outcomes were considered large (> 0.8). The physical activity levels of participants was not measured and this may have influenced the induction of cortical plasticity [9]. Finally, the influence of corticospinal potentiation on behavioural outcomes (e.g. motor skills, cognitive function, exercise tolerance etc.) and the relationship of this potentiation with GABA_A_ and GABA_B_ mediated inhibition [8, 12] remains unknown. These aspects will form an important extension of the current study.

### Conclusion & Significance

In conclusion, the study provides new evidence to show that ctDCS priming can improve the neuroplastic response (demonstrated with increased corticospinal excitability) to atDCS applied during low workload cycling exercise. Importantly, the findings of this study suggest that the typical lack of an increase in corticospinal response (known to represent activity-dependent neuroplasticity) seen post cycling exercise may be augmented with external stimuli (i.e. tDCS); and in particular, priming ctDCS optimizes the corticospinal tract for upcoming combined effect of cycling and atDCS. These preliminary findings are of high significance and will contribute to the development of tDCS priming protocols that may be used in sporting, clinical, rehabilitation and defence settings involving locomotor whole body movements to improve functional motor and cognitive function.

## Author Contributions

SS conceived and designed the study. MP and SS executed the study. MP and SS analysed data. MP and SS prepared figures and initial draft of the manuscript. SS and BL interpreted the data. All authors contributed to writing and approved the final version of the manuscript.

## Abbreviations

TMS: transcranial magnetic stimulation
MEPs: motor evoked potentials
atDCS: anodal transcranial direct current stimulation
ctDCS: cathodal transcranial direct current stimulation
M1: motor cortex
EMG: electromyography
FDI: first dorsal interosseous
RMT: resting motor threshold
NIBS: non-invasive brain stimulation

